# Convergent transcriptomic targets of propranolol and primidone identify potential biomarkers for essential tremor

**DOI:** 10.1101/2021.09.13.460137

**Authors:** Charles-Etienne Castonguay, Calwing Liao, Anouar Khayachi, Gabrielle Houle, Jay P Ross, Patrick A Dion, Guy A Rouleau

## Abstract

Essential tremor (ET) is one of the most common movement disorders, affecting nearly 5% of individuals over 65 years old. Despite its high heritability, few genetic risk loci for ET have been identified. Recent advances in pharmacogenomics have generated a wealth of data that led to the identification of molecular signatures in response to hundreds of chemical compounds. Among the different forms of data, gene expression has proven to be quite successful for the inference of drug response in cell models. We sought to leverage this approach in the context of ET where many patients are responsive two drugs: propranolol and primidone. Propranolol- and primidone-specific transcriptomic drug targets, as well as convergent gene targets across both drugs, could provide insights into the pathogenesis of ET and identify possible targets of interest for future treatments. In this study, cerebellar DAOY and neural progenitor cells were treated for 5 days with clinical concentrations of propranolol and primidone, after which RNA-sequencing was used to identify differentially expressed genes. The expression of genes previously implicated in genetic and transcriptomic studies of ET and other movement disorders, such as *TRAPPC11*, were significantly upregulated by propranolol. Pathway enrichment analysis identified multiple terms related to calcium signalling, endosomal sorting, axon guidance, and neuronal morphology. Convergent differentially expressed genes across all treatments and cell types were also found to be significantly more mutationally constrained, implying that they might harbour rare deleterious variants implicated in disease. Furthermore, these genes were enriched within cell types having high expression of ET related genes in both cortical and cerebellar tissues. Altogether, our results highlight potential cellular and molecular mechanisms associated with tremor reduction and identify relevant genetic biomarkers for drug-responsiveness in ET.

## INTRODUCTION

Essential tremor (ET) is one of the most common movement disorders^1^ affecting around 5% of individuals over 65 years old. The disease causes a 8-12 Hz kinetic tremor that typically affects the upper limbs but can also affect the head, voice, and rarely the lower limbs. Tremor intensity can sometimes increase with age and have a severe impact on activities of daily living. Recent studies aimed at identifying common and rare genetic variants have yielded mixed results, possibly due to clinical heterogeneity thus decreasing power of genetic studies^2^. Only a handful of variants have been identified and even fewer of them were replicated in other studies. Therefore, new approaches are needed, and transcriptomics might yield new insights in the pathophysiology of ET.

Recent studies in psychiatric genetics have successfully used drug effect screens to identify putative disease genes^3,4^. This approach is particularly relevant to diseases that have specific drug-responsive subsets of patients, as is the case with lithium responsive patients in bipolar disorder (BD)^5^. This kind of approach has yet to be used in many drug-responsive neurological disorders such as ET where patients respond to two drugs: propranolol and primidone^6^.

Propranolol and primidone are the most common drug treatments for ET. Both are efficient at reducing tremor by about 50% in ET patients^6^. Drug response is variable between patients, with some having a better outcome with either propranolol or primidone. Interestingly, some patients respond better to a combination of both drugs, especially for reducing limb and head tremors, hinting at potential additive or synergistic effects^7^. Propranolol is a beta-adrenergic receptor 1/2 antagonist initially developed to treat hypertension. In the context of ET, propranolol is thought to act on peripheral beta-2 receptors in muscle spindles, but it also has effects on cells in the central nervous system (CNS)^7,8^. Propranolol is lipophilic enough to cross the blood brain barrier (BBB) and to accumulate in high concentrations in mouse cerebellum and cortex following treatment^9^. Primidone is an anticonvulsant whose mechanism of action in ET is not well defined but it possibly reduces calcium and sodium currents across neuronal membranes^6^, therefore, reducing neuronal excitability.

The transcriptomic effects of primidone and propranolol in the context of ET remain poorly understood^10,11^. Propranolol increased the expression *SHF*, a gene that was shown to be downregulated in ET patient cerebellum^10^. Studying the effects of tremor-reducing drugs on transcription can inform us on mechanisms that reduce tremors. Furthermore, it is possible that genes that are targeted by both drugs are implicated in ET pathophysiology and could allow for the identification of genes harbouring putative ET causing variants.

In this study, we identified convergent transcriptomic targets of primidone and propranolol in cortical neural progenitor cells (NPC) and cerebellar medulloblastoma cells (DAOY). Common cellular pathways affected by both treatments were related to neuronal morphology, axon guidance as well as cell-cell interactions as revealed by co-expression and pathway enrichment analysis. We also found that ET drugs specifically affected the expression of genes intolerant to loss-of-function (LoF) variants, hinting at possible enrichment of such rare LoF variants. Furthermore, with integration of single-cell data, we find that drug-targeted genes are mostly enriched in non-neuronal cell types such as endocytes, astrocytes, and oligodendrocytes in both cortical and cerebellar tissues. Our study identifies new putative ET- and tremor-related genes and informs on the molecular and cellular basis for tremor-reduction in ET.

## METHODS

### Cell culture and drug treatment

DAOY and NPC cells were cultured as previously described^5,11^ and treated for 5 days with 20 ng/mL of propranolol or 5 μg/mL of primidone (n = 3 per treatment/cell line). H2O- or DMSO (0.023%)-treated cells were used as controls for propranolol and primidone, respectively. Drug concentrations were chosen based on previous studies that tested efficient tremor-reducing serum levels of propranolol and primidone in ET patients^12,13^. A kill curve was used to determine lethal drug concentrations for DAOY cells and NPCs in culture (Supplementary Table 10-12, Supplementary Figure 1-2).

### RNA-sequencing and differential expression analysis

RNA was extracted with the RNeasy Mini Kit (Qiagen). cDNA library preparation was done using NEBNext stranded library preparation protocol (New England Biolabs) with rRNA depletion using the QIAseq FastSelect rRNA HMR kit. (Qiagen). Samples were sequenced on the Illumina NovaSeq6000 platform (150bp paired-end reads, 150M reads). FASTQ files were pseudo-aligned to the Ensembl v102 annotation of the human genome using Salmon v1.4.0^14^. Gene-level differential expression analysis was done using the R package Sleuth^15^. Only genes with a minimum of 10 scaled reads per base in 90% of samples were kept to filter out low-count genes. Cell types and treatments were analyzed separately using the Wald test (WT). The full model for the WT was:

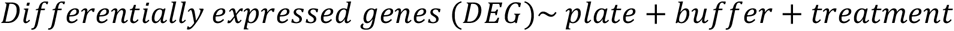

MA plots and p-value histograms displayed expected distributions (Supplementary Figure 3,4). Meta-analysis of gene Z-scores was performed to analyze convergent DEG across cell types and treatments. Briefly, Z-scores for each gene were calculated and then summed across different combinations of cell types and treatments using Stouffer’s Z method^16^. Multiple analyses were performed notably propranolol specific effect across cell types (labeled ‘prop’; Supplementary table 6), primidone effect across cell types (‘prim’; Supplementary table 7), convergent propranolol and primidone effect in each cell type (‘daoy’ and ‘npc’; Supplementary tables 8 and 9 respectively) and convergent primidone and propranolol effects across both cell types (‘all’; Supplementary table 5). False discovery rate was controlled for using the Benjamini-Hochberg procedure (q-value threshold < 0.05). At least 3 DEGs with highest fold-change per condition were validated using TaqMan qPCR probes (Supplementary Table 13).

### WGCNA, co-expression and pathway enrichment

WGCNA was done using the R package^17^. DAOY and NPC sequencing results were analyzed separately, merging both primidone and propranolol treatments in the analysis. Normalized TPM values obtained from Sleuth (‘sleuth_to_matrix’) were used for the analysis. To filter out noisy low-count genes, only genes with a minimum of 10 TPM in 47% of samples were kept, for a final list of 8549 genes in DAOYs and 9260 genes for NPC. Two outlier samples (‘DAOY_PRIM_03’ and ‘NPC_PRIM_02’) were removed from the analysis based on sample clustering dendrogram. Fisher’s exact test was used to calculate gene-module p-values. Co-expression analysis was performed using GeneNetwork2.0^18^. Pathway enrichment analysis was done using the gprofileR R package^19^. Briefly, gene-lists were made from convergent DEGs across multiple conditions (both drugs in DAOYs or NPCs, propranolol or primidone in both cells, both drugs in both cells). Custom background used in gprofiler comprised genes expressed in either DAOYs, NPCs or both when pertinent. The g:SCS algorithm was used for multiple testing correction (q-value threshold < 0.1).

### Correlation with ET TWAS summary statistics

ET TWAS summary statistics were obtained from Liao et al. (2021; unpublished results). A generalized linear model was used to measure the strength of association between gene-level drug Z-scores and TWAS Z-scores, controlling for gene length and gene GC content (‘lm’ function in R). Weighted Z-scores were also used to account for significance of effect. The formula used were:

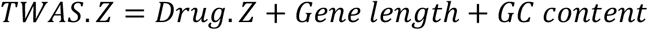

And for the weighted Z-score analysis:

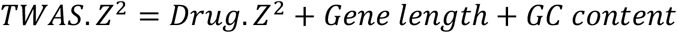

Association p-values were corrected for multiple testing using Benjamini-Hochberg (q-value threshold < 0.05).

### Single cell enrichment analysis

A one sample Z-test was used to test enrichment of drug-targeted genes as described previously^20^. An ET gene-set was curated from genes associated with ET from linkage, whole-exome, GWAS and transcriptomic studies ^2,10^. Drug gene-sets were made from convergent DEGs (FDR < 0.05) across different conditions (DAOY, NPC, propranolol, primidone, all conditions). Adult cerebellum single-nucleus RNA sequencing data was obtained from Lake et al. (2018; GEO accession: <u>GSE97930</u>)^21^. Average cell counts per cell-type were obtained using Seurat v4.0.1^22^. Trimmed means per cell-type from adult cortex single-cell RNA-sequencing were obtained from the Allen Brain Atlas Smart-seq multiple cortical regions dataset^23^. To account for drop-out rates and reduce zero-inflation of the single-cell count matrices, low average count genes were filtered out in both cerebellum (< 0.5 counts in 7/10 cell types) and cortex (<1 count in 85/121 cell types). Single sample Z-tests were used to obtain cell-type specific enrichment Z-scores:

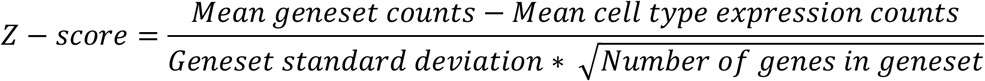

### Loss-of-function analysis

The distribution of mutational constraint scores for drug DEGs was assessed using pLoF o/e ratio scores obtained from gnomAD^24^. pLof scores for convergent genes across all conditions with q-val <0.05 were compared all protein coding genes passing QC from the Sleuth differential expression analysis. To account for coding sequence length and gene GC percentage, propensity score matching with replacement was used (matchIT package in R^25^) to measure pLoF score distribution differences between DE drug genes and all protein coding genes included in the meta-analysis. Nearest neighbor matching with the maximum number of matches (ratio = 1:43) between non-DEGs and DEGs was used. A Wilcoxon unpaired test was done on the matched data. The same methods were used to assess pLoF score differences of upregulated (match ratio = 1:57) and downregulated (match ratio = 1:178) DEGs with all protein coding genes.

## RESULTS

### Differential expression following propranolol and primidone treatment

To assess the transcriptomic effect of propranolol and primidone on neuronal and cerebellar cells, NPCs and DAOYs were independently treated with clinically relevant concentrations of both drugs for five days. Treatment of DAOYs with propranolol resulted in 1,754 DE genes (Supplementary Table 1) while treatment of NPCs resulted in 1,571 DE genes (Supplementary Table 2). Directionality of overall transcriptional effect was widely different between NPCs and DAOYs, with propranolol treatment resulting in mostly overexpression in DAOYs and underexpression in NPCs (Figure 1C and 1D). Pearson correlation of propranolol-treated NPCs and DAOYs effectively show a strong negative correlation, indicating opposite transcriptomic effects on the same genes (r = -0.35, p-val < 2.2E-308, Figure 1A). However, this correlation weakens when weighing for the most significant DEGs (r = -0.283, p = 7.1E-214, Figure 1B). Primidone, on the other hand, had a weak effect on transcription in both NPCs and DAOYs with only 200 (Supplementary table 4) and 23 DEGs (Supplementary table 3) in each, respectively. In NPCs, propranolol and primidone DEGs were lowly correlated (r = -0.06, p-val = 1.6E-11, Figure 1A) with a weaker weighted correlation (r = -0.021, p-val = 2.2E-02, Figure 1B). Similar weak (weighted and unweighted) correlations are seen between the two drugs in DAOYs (Figure 1A and 1B).

**Figure 1.**
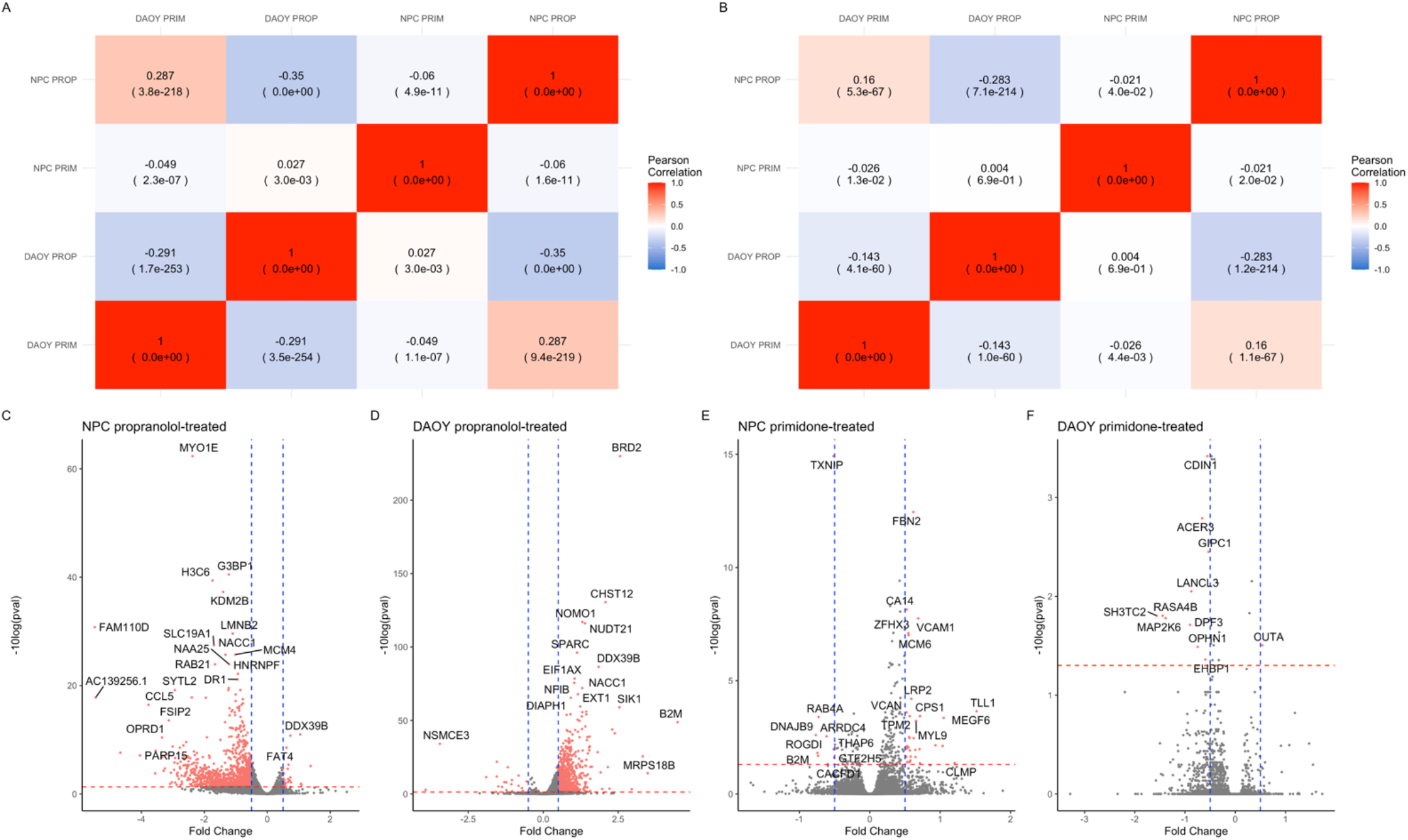
Correlation between DAOYs and NPCs treated with propranolol and primidone. A. Unweighted Pearson correlations between DEGs z-scores from different conditions of treatment and cell types. B. Weighted Pearson correlations between DEGs z-scores from different conditions of treatment and cell types. C-F. Volcano plots of propranolol-treated NPCs (C) and DAOYs (D) as well as primidone-treated NPCs (E) and DAOYs (F). Blue lines indicate -0.5- and 0.5-fold changes. Red lines indicate q-value significance threshold (0.05).

### ET drug targets converge on genes related to movements disorders and ET

Shared effects of propranolol and primidone on specific genes increases the likelihood of these genes being integral to tremor reduction in ET. Therefore, convergence of drug effects on expression was assessed by comparing gene Z-scores from different treatment conditions: convergent drug targets in either DAOYs or NPCs, convergent propranolol or primidone targets in both cell types and convergent targets of both drugs in all cell types.

Across DAOYs and NPCs, 788 significant convergent DEGs were found with propranolol treatment (Supplementary table 6) and 36 convergent DEGs following primidone treatment (Supplementary table 7). Propranolol, in both cell types, increased expression of *TRAPPC11*, a trafficking protein previously associated with ET^26^ (z-score = 5.41, p-val = 5.87E-06). Propranolol also decreased expression of *G3BP1* (z =-9.07, q-val = 7.84E-17), which encodes a protein implicated in stress granule formation and is known to affect axonal mRNA translation as well as nerve regeneration^27^. *BRD2*, a transcription factor previously implicated with epilepsy, was upregulated following propranolol treatment in both cells (z = 21.13, q-val = 4.56E-95). *NONO (*z *=* 6.93, q-val = 3.69E-09), a gene harbouring a splicing variant known to cause X-linked intellectual deficiency with intentional tremor, was found to be upregulated^28^. Primidone, across NPCs and DAOYs, upregulated *VCAM1 (*z-score = 5.53, p-value = 1.29E-04), a gene implicated in axonal myelination by oligodendrocytes^29^. *GIPC1* was also found to be downregulated following primidone treatment in both cell types (z = -5.46, q-val = 1.42E-04). GIPC1 is a known interactor of DRD3 which has previously been associated with ET and Parkinson’s (PD)^2,30,31^.

### Propranolol and primidone act on pathways related to neuronal survival as well as axon guidance

Following the identification of convergent DEGs across treatments, we wanted to identify molecular pathways affected by propranolol and primidone in DAOYs and NPCs. Co-expression enrichment analysis (using GeneNetwork2.0^18^) for convergent DEGs across all conditions showed that Reactome terms related to GPCR signalling (p-val = 1.12E-19), axon guidance (p-val = 1.68E-08), Semaphorin interactions (p-val = 3.24E-13) and VEGF signalling (p-val = 2.23E-08) were significantly enriched within the convergent genesets (Supplementary Table 14). Furthermore, Ca^2+^ signalling (p-val = 4.67E-07) and voltage-gated potassium channels (p-val = 4.64E-06) were also found to be significantly enriched. Interestingly, GO:cellular components significant terms were mostly related to cell:cell or cell:extracellular matrix interactions as well as axon guidance such as lamellipodium (p-val = 4.47E-13), filopodium (p-val = 3.54E-11, focal adhesion (p-val = 4.70E-11) and growth cone (p-val = 1.04E-09)(Supplementary table 16).

Pathway enrichment analysis of convergent propranolol DEGs (in both cell types) was also performed using g:profiler using genes expressed in both DAOYs and NPCs as background (Table 1). Pathways known to be affected by propranolol such as HIF-1*α (*p-val = 0.001) and regulation of apoptosis (p-val = 0.02) were significantly enriched. Much like the co-expression analysis, Reactome terms related to axon guidance were found to be significant, such as RUNX1 transcription (p-val = 0.0002), a transcription factor implicated in growth cone guidance of DRG neurons^32^. Interestingly, CaMKK2 signalling pathway was found to be significantly enriched within genes in the propranolol geneset. *CAMKK2* encodes a kinase implicated in synapse homeostasis and is also involved in modyifing Aβ synaptotoxicity in Alzheimer’s disease^33^.

**Table 1.** Pathway enrichment for convergent propranolol DEGs in both DAOYs and NPCs.

| SOURCE | TERM | P-VALUE |
| --- | --- | --- |
| CORUM | PA700 complex | 0.00732592 |
| CORUM | p54(nrb)-PSF-matrin3 complex | 0.00741609 |
| CORUM | PA700-20S-PA28 complex | 0.01284008 |
| CORUM | HEXIM1-DNA-PK-paraspeckle components-<br>ribonucleoprotein complex | 0.05052404 |
| CORUM | Ubiquitin E3 ligase (CHEK1, CUL4A) | 0.06576926 |
| CORUM | CORUM root | 0.07664168 |
| CORUM | EBAFb complex | 0.08852844 |
| CORUM | NCOR1 complex | 0.08852844 |
| KEGG | Proteasome | 0.00921554 |
| KEGG | Spinocerebellar ataxia | 0.02672326 |
| KEGG | Prion disease | 0.04664458 |
| **KEGG** | Protein processing in endoplasmic reticulum | 0.05311146 |
| **KEGG** | Hippo signaling pathway - multiple species | 0.08972819 |
| **MIRNA** | hsa-miR-6766-5p | 0.00036715 |
| **MIRNA** | hsa-miR-6756-5p | 0.00036715 |
| **MIRNA** | hsa-miR-539-5p | 0.0003869 |
| **MIRNA** | hsa-miR-4668-3p | 0.00716318 |
| **MIRNA** | hsa-miR-21-5p | 0.0132699 |
| **MIRNA** | hsa-miR-654-5p | 0.02081865 |
| **MIRNA** | hsa-miR-541-3p | 0.02687402 |
| **MIRNA** | hsa-miR-1468-3p | 0.0441487 |
| **MIRNA** | hsa-let-7b-5p | 0.04603661 |
| **MIRNA** | hsa-miR-548f-5p | 0.05118884 |
| **MIRNA** | hsa-miR-548aj-5p | 0.05470749 |
| **MIRNA** | hsa-miR-548x-5p | 0.05470749 |
| **MIRNA** | hsa-miR-548g-5p | 0.05470749 |
| **MIRNA** | hsa-miR-193b-3p | 0.05509061 |
| **REAC** | Transcriptional regulation by RUNX1 | 0.00022561 |
| **REAC** | Oxygen-dependent proline hydroxylation of Hypoxia-inducible Factor Alpha | 0.0011874 |
| **REAC** | Cellular response to hypoxia | 0.00350166 |
| **REAC** | Host Interactions of HIV factors | 0.00421087 |
| **REAC** | Cell Cycle Checkpoints | 0.00665026 |
| **REAC** | UCH proteinases | 0.007029 |
| **REAC** | G2/M Checkpoints | 0.01195953 |
| **REAC** | Regulation of ornithine decarboxylase (ODC) | 0.01244161 |
| **REAC** | G1/S DNA Damage Checkpoints | 0.01314543 |
| **REAC** | Signaling by NOTCH | 0.01416007 |
| **REAC** | p53-Independent G1/S DNA damage checkpoint | 0.01463202 |
| **REAC** | Ubiquitin Mediated Degradation of Phosphorylated Cdc25A | 0.01463202 |
| **REAC** | p53-Independent DNA Damage Response | 0.01463202 |
| **REAC** | Regulation of APC/C activators between G1/S and early anaphase | 0.0153801 |
| **REAC** | Regulation of Apoptosis | 0.01714052 |
| **REAC** | Cdc20:Phospho-APC/C mediated degradation of Cyclin A | 0.02113246 |
| **REAC** | Assembly of the pre-replicative complex | 0.02316267 |
| **REAC** | Deubiquitination | 0.02357632 |
| **REAC** | Autodegradation of Cdh1 by Cdh1:APC/C | 0.02437405 |
| **REAC** | APC:Cdc20 mediated degradation of cell cycle proteins prior to satisfaction of the cell cycle checkpoint | 0.02451223 |
| **REAC** | Regulation of MECP2 expression and activity | 0.02941481 |
| **REAC** | Stabilization of p53 | 0.03112835 |
| REAC | APC/C:Cdc20 mediated degradation of mitotic proteins | 0.03270423 |
| REAC | DNA Replication Pre-Initiation | 0.03291172 |
| REAC | Orc1 removal from chromatin | 0.03381524 |
| REAC | PTEN Regulation | 0.03447437 |
| REAC | Metabolism of polyamines | 0.03559536 |
| REAC | Activation of APC/C and APC/C:Cdc20 mediated degradation of mitotic proteins | 0.03762377 |
| REAC | Regulation of mitotic cell cycle | 0.03959616 |
| REAC | APC/C-mediated degradation of cell cycle proteins | 0.03959616 |
| REAC | Transcriptional regulation by RUNX3 | 0.03975456 |
| REAC | CDT1 association with the CDC6:ORC:origin complex | 0.04112471 |
| REAC | MAPK6/MAPK4 signaling | 0.04224034 |
| REAC | Ub-specific processing proteases | 0.04291446 |
| REAC | Switching of origins to a post-replicative state | 0.04311326 |
| REAC | APC/C:Cdc20 mediated degradation of Securin | 0.045216 |
| REAC | Vpu mediated degradation of CD4 | 0.05357088 |
| REAC | Cross-presentation of soluble exogenous antigens (endosomes) | 0.07194281 |
| REAC | Regulation of activated PAK-2p34 by proteasome mediated degradation | 0.07194281 |
| REAC | Hedgehog ligand biogenesis | 0.074096 |
| REAC | p53-Dependent G1/S DNA damage checkpoint | 0.08163726 |
| REAC | p53-Dependent G1 DNA Damage Response | 0.08163726 |
| REAC | SCF-beta-TrCP mediated degradation of Emi1 | 0.0874049 |
| REAC | CDK-mediated phosphorylation and removal of Cdc6 | 0.09137813 |
| REAC | Autodegradation of the E3 ubiquitin ligase COP1 | 0.09544567 |
| REAC | Ubiquitin-dependent degradation of Cyclin D | 0.09544567 |
| WP | mRNA Processing | 0.00409008 |
| WP | CAMKK2 Pathway | 0.00436354 |
| WP | Pathways Affected in Adenoid Cystic Carcinoma | 0.01716516 |
| WP | MET in type 1 papillary renal cell carcinoma | 0.02394081 |
| WP | Oncostatin M Signaling Pathway | 0.07825036 |
| WP | 15q13.3 copy number variation syndrome | 0.07966433 |
| WP | Gastrin Signaling Pathway | 0.09031422 |

Weighted gene correlation network analysis was also performed to identify co-expression modules associated with combined propranolol/primidone treatment. Module-trait and module correlation heatmaps are shown in Figure 2. Two modules (cyan and red; corr = 0.74, p-val = 0.009; corr = 0.73, p-val = 0.01 respectively; Figure 2A) were found to be significantly associated with treatment in DAOYs and only one module (red; corr = 0.65, p-val = 0.03) was significantly associated with NPCs (Figure 2B). Pathway enrichment analysis of DAOY red module genes found an enrichment of Reactome terms related to RABGAP signalling (p-val = 0.009) as well as RUNX1 transcription (p-val = 0.02; Table 2). NPC red modules genes were significantly associated with neuronal morphology, axon guidance and neurogenesis (Table 3).

**Figure 2.**
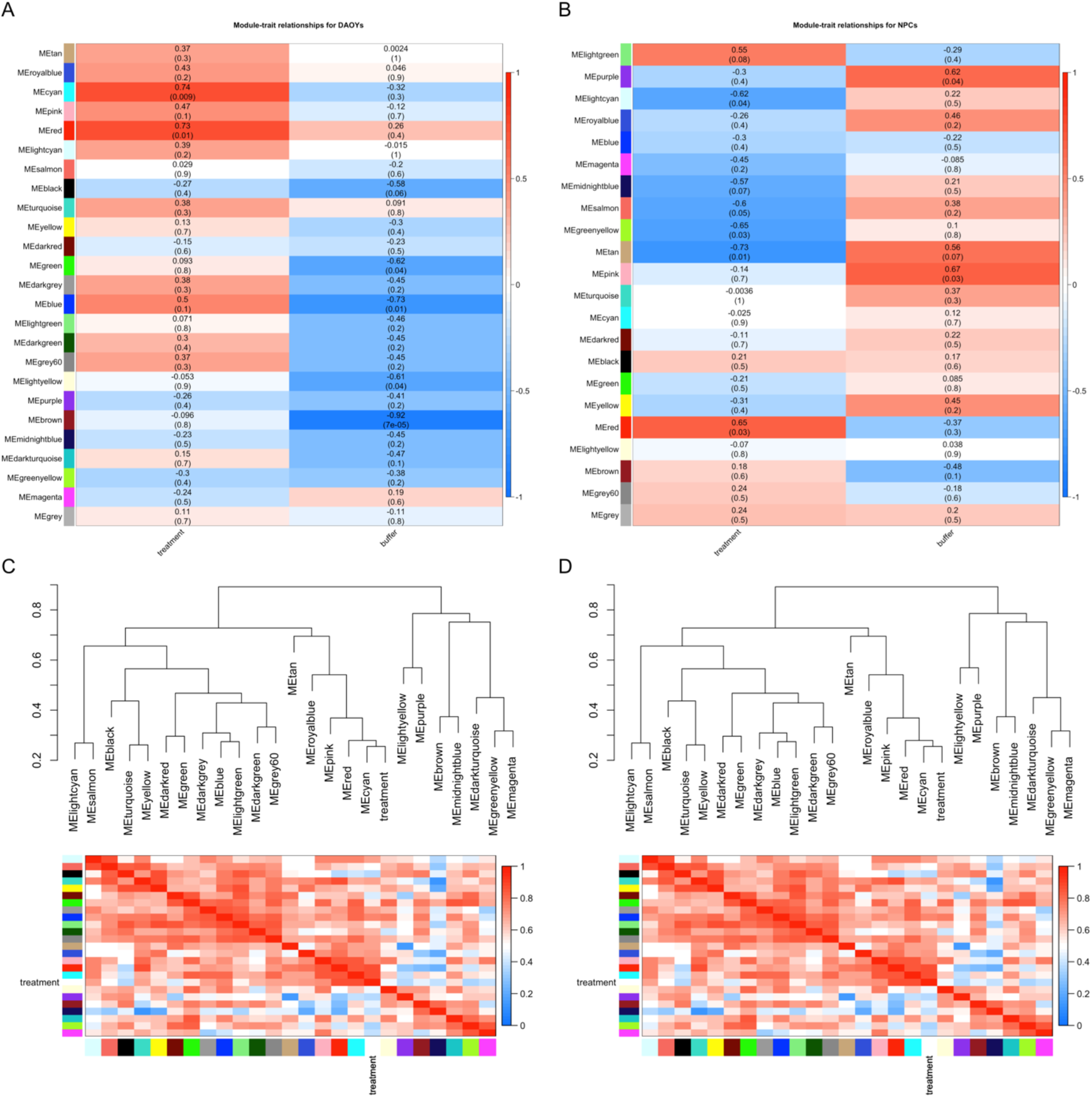
Co-expression gene modules for convergent propranolol and primidone targets. A. Module-treatment (propranolol/primidone) and -buffer (H2O/DMSO; control) correlation heatmaps for DAOYs. B. Module-treatment (propranolol/primidone) and -buffer (H2O/DMSO; control) correlation heatmaps for NPCs. Value indicates correlation between gene-trait and gene-module associations with p-value in parenthesis. C. Module dendrograms with module membership correlation heatmaps for DAOYs. D. Module dendrograms with module membership correlation heatmaps for NPCs.

**Table 2.** Pathway enrichment analysis of red gene module for drug treatment in DAOYs.

| SOURCE | TERM_NAME | P_VALUE |
| --- | --- | --- |
| CORUM | Ubiquitin E3 ligase (CCDC22, COMMD8, CUL3) | 0.00491141 |
| **CORUM** | Ecsit complex (ECSIT, MT-CO2, GAPDH, TRAF6, NDUFAF1) | 0.07383335 |
| **REAC** | TBC/RABGAPs | 0.00987381 |
| **REAC** | RUNX3 regulates YAP1-mediated transcription | 0.02324914 |
| **REAC** | RNA polymerase II transcribes snRNA genes | 0.08552303 |
| **REAC** | Rab regulation of trafficking | 0.09310043 |
| **WP** | Eukaryotic Transcription Initiation | 0.09003334 |

**Table 3.**
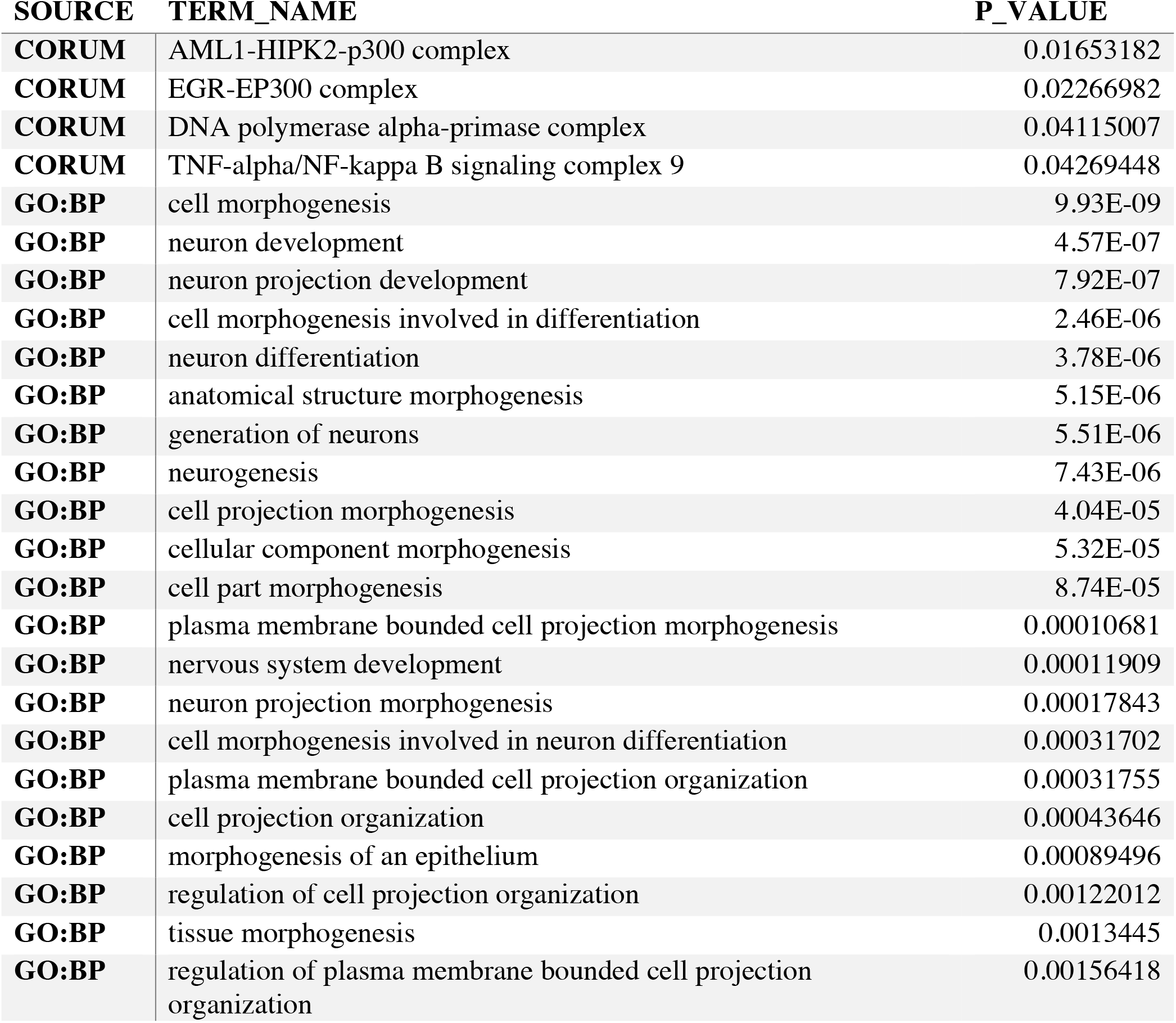
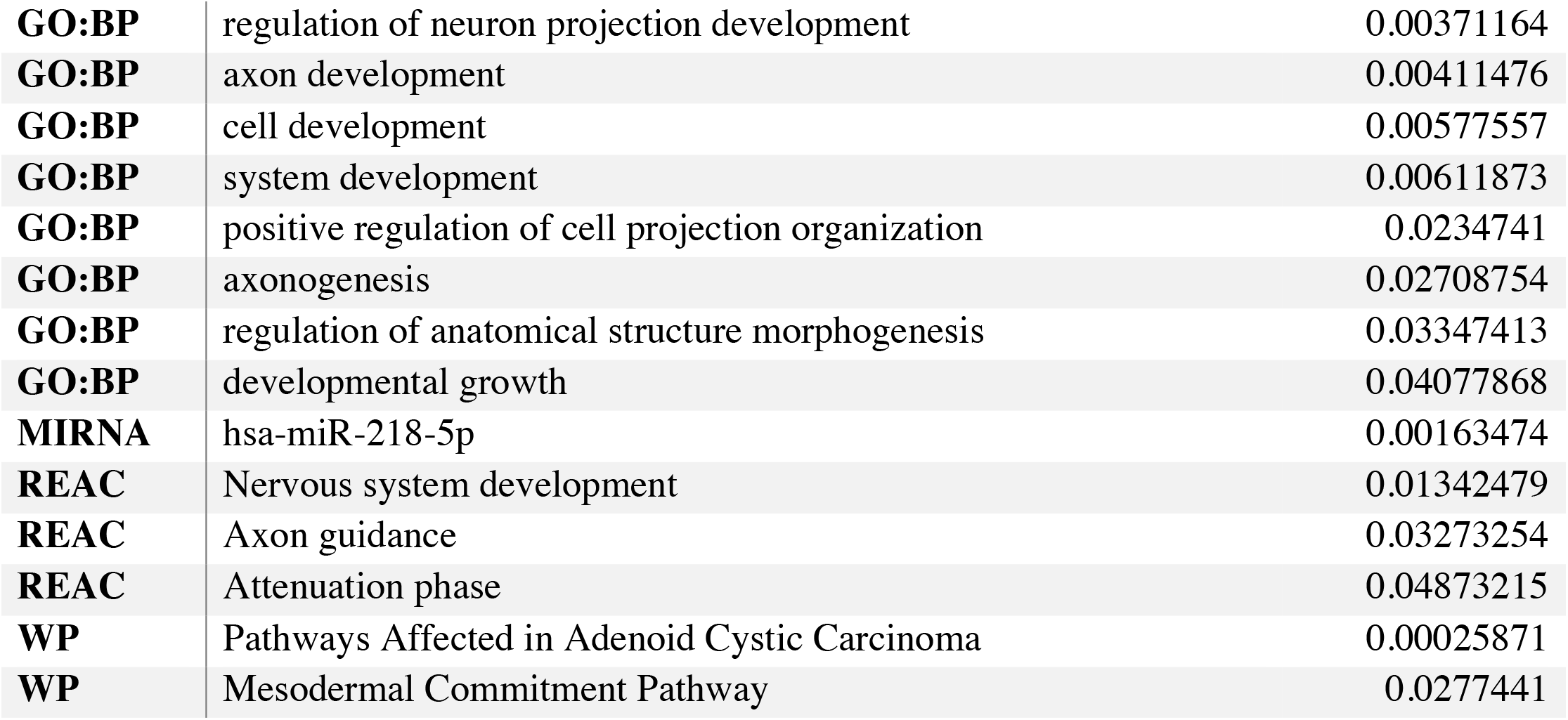
Pathway enrichment analysis of red gene module for drug treatment in NPCs.

| **SOURCE** | **TERM_NAME** | **P_VALUE** |
| --- | --- | --- |
| **CORUM** | AML1-HIPK2-p300 complex | 0.01653182 |
| **CORUM** | EGR-EP300 complex | 0.02266982 |
| **CORUM** | DNA polymerase alpha-primase complex | 0.04115007 |
| **CORUM** | TNF-alpha/NF-kappa B signaling complex 9 | 0.04269448 |
| **GO:BP** | cell morphogenesis | 9.93E-09 |
| **GO:BP** | neuron development | 4.57E-07 |
| **GO:BP** | neuron projection development | 7.92E-07 |
| **GO:BP** | cell morphogenesis involved in differentiation | 2.46E-06 |
| **GO:BP** | neuron differentiation | 3.78E-06 |
| **GO:BP** | anatomical structure morphogenesis | 5.15E-06 |
| **GO:BP** | generation of neurons | 5.51E-06 |
| **GO:BP** | neurogenesis | 7.43E-06 |
| **GO:BP** | cell projection morphogenesis | 4.04E-05 |
| **GO:BP** | cellular component morphogenesis | 5.32E-05 |
| **GO:BP** | cell part morphogenesis | 8.74E-05 |
| **GO:BP** | plasma membrane bounded cell projection morphogenesis | 0.00010681 |
| **GO:BP** | nervous system development | 0.00011909 |
| **GO:BP** | neuron projection morphogenesis | 0.00017843 |
| **GO:BP** | cell morphogenesis involved in neuron differentiation | 0.00031702 |
| **GO:BP** | plasma membrane bounded cell projection organization | 0.00031755 |
| **GO:BP** | cell projection organization | 0.00043646 |
| **GO:BP** | morphogenesis of an epithelium | 0.00089496 |
| **GO:BP** | regulation of cell projection organization | 0.00122012 |
| **GO:BP** | tissue morphogenesis | 0.0013445 |
| **GO:BP** | regulation of plasma membrane bounded cell projection organization | 0.00156418 |
| **GO:BP** | regulation of neuron projection development | 0.00371164 |
| **GO:BP** | axon development | 0.00411476 |
| **GO:BP** | cell development | 0.00577557 |
| **GO:BP** | system development | 0.00611873 |
| **GO:BP** | positive regulation of cell projection organization | 0.0234741 |
| **GO:BP** | axonogenesis | 0.02708754 |
| **GO:BP** | regulation of anatomical structure morphogenesis | 0.03347413 |
| **GO:BP** | developmental growth | 0.04077868 |
| **MIRNA** | hsa-miR-218-5p | 0.00163474 |
| **REAC** | Nervous system development | 0.01342479 |
| **REAC** | Axon guidance | 0.03273254 |
| **REAC** | Attenuation phase | 0.04873215 |
| **WP** | Pathways Affected in Adenoid Cystic Carcinoma | 0.00025871 |
| **WP** | Mesodermal Commitment Pathway | 0.0277441 |

### Correlation of the effects of propranolol and primidone with those of common and rare variants in ET

TWAS studies the effect of common SNPs associated with a disease on the expression of genes in different tissues. We postulated that transcriptomic targets of propranolol and primidone might correlate with the transcriptomic effect of common ET variants. We used TWAS summary statistic from an upcoming ET GWAS (Liao et al., unpublished results) to measure the correlation between TWAS gene Z-scores and convergent drug target Z-scores (across all possible conditions) while controlling for gene length and GC content. Weak, non-significant correlations between TWAS Z-scores and drug target Z-scores were found across all conditions and all brain tissues (p > 0.05; Figure 3A). Cerebellar hemispheres and cerebellum tissues, brain regions highly associated with ET, displayed non-significant negative correlations with convergent drug targets (coeff = -0.0143, p-val = 0.549; coeff = -0.000138, p-val = 0.994 respectively; Figure 3B).

**Figure 3.**
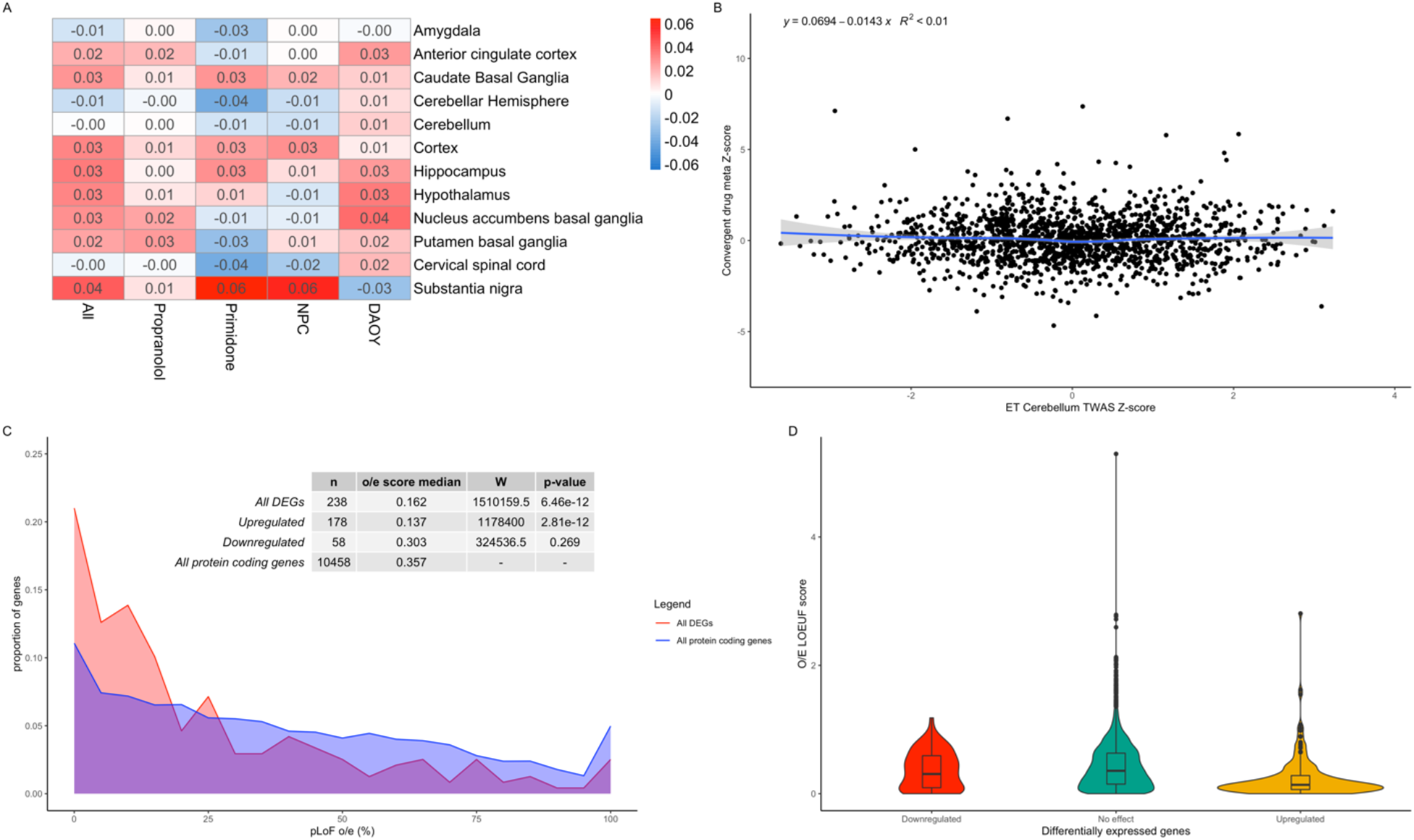
Effects of ET drugs on common and rare variants. A. Correlation heatmap of ET TWAS gene Z-scores in different brain tissues and drug effect gene Z-scores from different meta-analysis conditions. Values indicate Z-score regression coefficient from linear model. B. Correlation plot of TWAS gene Z-scores from cerebellar tissue and convergent primidone and propranolol gene Z-scores across DAOYs and NPCs. C. Line histogram displaying the distribution of O/E LOEUF scores from all protein coding genes (blue) and convergent DEGs (red) following drug treatment. O/E scores were directly transformed to percentages (ex. 0.25 as 25%) with scores over 10 counted as 100%. D. Violin plots of O/E LOEUF scores for upregulated DEGs (yellow), dowregulated DEGs (red) and non-significant DEGs (green).

We postulated that since propranolol and primidone had a non-significant correlation with expression of genes harbouring common variants for ET, they might instead act on genes that have rare variants. GnomAD recently published observed/expected (o/e) loss-of-function (LoF) scores for all protein coding genes in the genome. These scores inform on the tolerance of genes to rare LoF variants, with genes with a higher frequency of observed to expected LoF variants being more tolerant to mutations. Figure 3C shows the distribution of LoF scores of drug DEGs compared to all protein coding genes passing the initial DE QC. Drug targets displayed a significantly lower o/e score median (n = 256, median = 0.18) than all protein coding genes (n = 11,188, median = 0.36; W = 1727520, p-val = 1.501E-10) using a Wilcoxon unpaired test. Interestingly, when looking at fold change direction (figure 3D), upregulated genes (n = 194) had a significantly lower o/e score median (median = 0.15, W = 1361482, p-value = 2.917E-12) than all protein coding genes whilst no significant difference was found between o/e scores medians of downregulated genes (n = 71) and all protein coding genes (median = 0.35, W = 417126, p-value = 0.3246) using a Wilcoxon unpaired test. Thus, propranolol and primidone increased expression of mutationally constrained genes in cultured DAOYs and NPCs.

### Single cell enrichment of propranolol and primidone targeted genes

Our current understanding of CNS cell types affected in ET is still very limited. Enrichment of disease related genes can indirectly inform on potential cell types implicated in disease pathophysiology^20^. We first sought to assess the enrichment of ET genes discovered through familial linkage studies as well as whole-exome studies in cell types of the adult cerebellum and cerebral cortex (Figure 4, Supplementary Table 17-18). Enrichment Z-scores per cell type for ET genes as well as drug DEGs were calculated based on average normalized expression in single nucleus cerebellum data from Lake et al. (2018)^21^ and cortical single-cell Smart-seq data from the Allen Brain Institute. In the cerebellum, ET genes were mostly enriched in astrocytes (enrichment z-score = 3.11, q-value = 0.021; Figure 4A and 4B). In the cortex, the strongest enrichments of ET genes were found in oligodendrocyte progenitor cells (OPCs) (z-score = 3.55) and L3-L5 excitatory neurons with the most significant neuronal cell type being the *FEZF2-, DYRK*-expressing pyramidal neurons of cortical layer V (z-score = 3.28, q-val = 0.0068; Figure 4C). Significant enrichment was also found in L1 *MTG1* astrocytes (z-score = 3.13, q-val = 0.0090).

**Figure 4.**
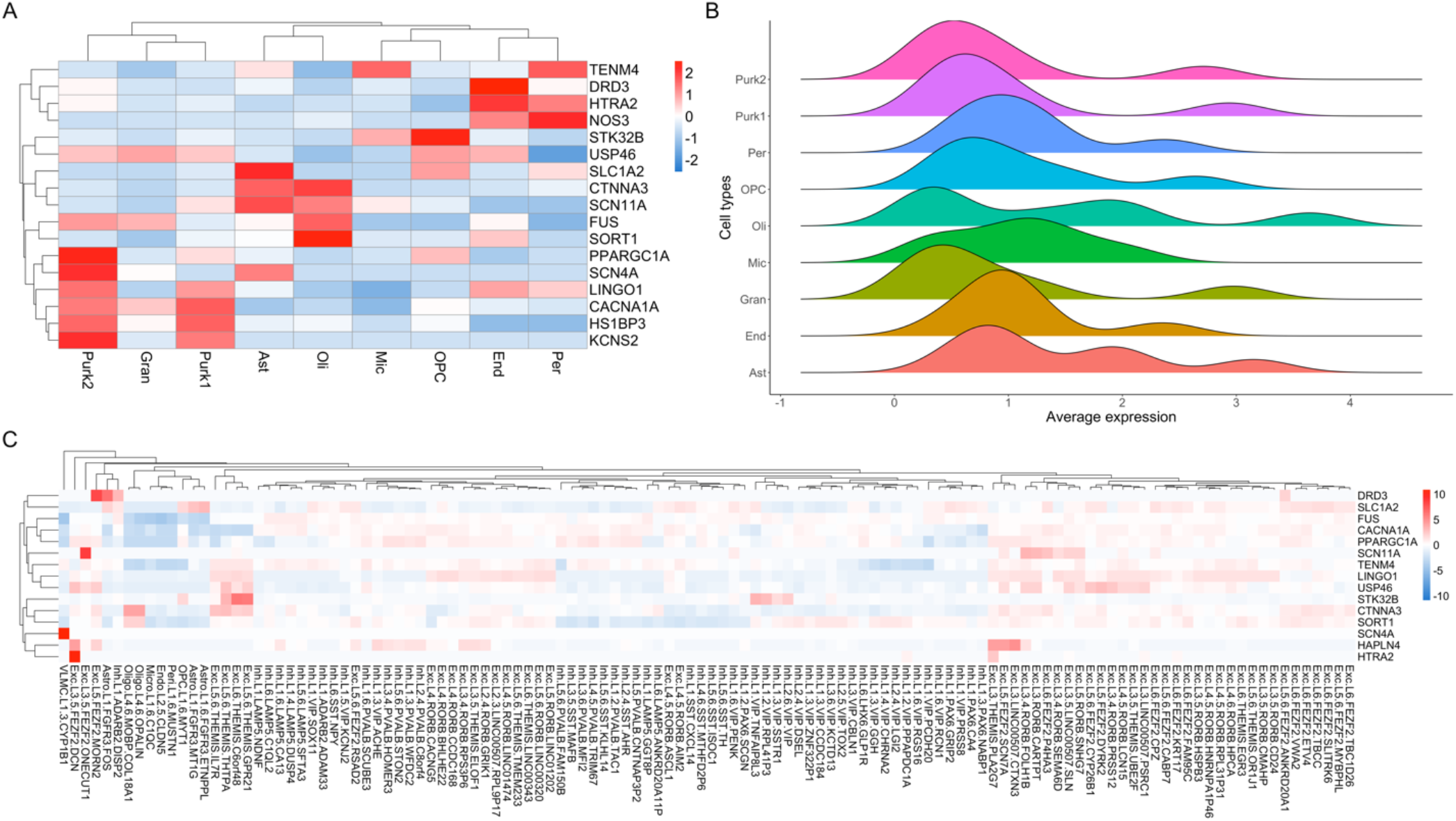
Single-cell enrichment of ET genes in cerebellar and cortical tissues. A. Single-cell enrichment Z-score heatmap of ET-related genes in adult cerebellar tissue. Rows represent ET genes; Columns represent cell types of the cerebellum (Purk1 = SORC3+ Purkinje cells, Purk2 = SORC3-Purkinje cells, Ast = Astrocytes, OPC = Oligodendrocyte progenitor cells, Oli = Oligodendrocytes, Mic =Microglia, End = Endocytes, Gran = Granule cells, Per = Pericytes). B. Ridge plots displaying distribution of average expression counts of ET-related genes in different cell types of the adult cerebellum. C. Z-score expression heatmap of ET genes in single-cell types of the adult cortex. Rows represent ET genes; Columns represent cortical cell types (Exc = Excitatory, Inh = Inhibitory, L# = cortical layer, Astro = Astrocytes).

Next, we assessed the enrichment of propranolol and primidone DEGs identified in this study in cortical and cerebellar single-cell data (Figure 5, Supplementary Table 17-18). In cerebellum single-nucleus data, convergent propranolol DEGs were mostly enriched in endocytes (z-score = 3.38, q-val = 0.014) and microglia (z-score = 3.36, q-val = 0.014) whilst convergent propranolol/primidone DEGs in all cell types were mostly enriched in oligodendrocytes (z-score = 2.90, q-val = 0.034; Figure 5E). Interestingly, convergent propranolol/primidone DEGs in DAOYs, a cell-type specific to the cerebellum, had enriched expression in astrocytes (z-score = 2.74, q-val = 0.047), much like the enrichment of ET genes in cerebellar astrocytes (Figure 4A). In cortical tissue, convergent drug DEGs were mostly significantly enriched in non-neuronal cell types (figure 5D), notably oligodendrocytes (z-score = 5.09, q-val = 3.65E-07), astrocytes (z-score. = 4.92, q-val = 1.00E-04) and endocytes (z-score = 3.95, q-val = 1.70E-03). Unsurprisingly, given the use of propranolol to lower blood pressure, convergent propranolol DEGs were mostly enriched in endocytes (z-score = 6.18, q-val = 4.48-07) and vascular and leptomeningeal cells (VLMC; z-score = 4.77, q-val = 1.52E-04). Of note, propranolol DEGs were also enriched in L1-L3 inhibitory neurons, notably vasoactive intestinal peptide (VIP) expressing inhibitory neurons (Figure 5D, see Supplementary Table 17 and 18 for statistics).

**Figure 5.**
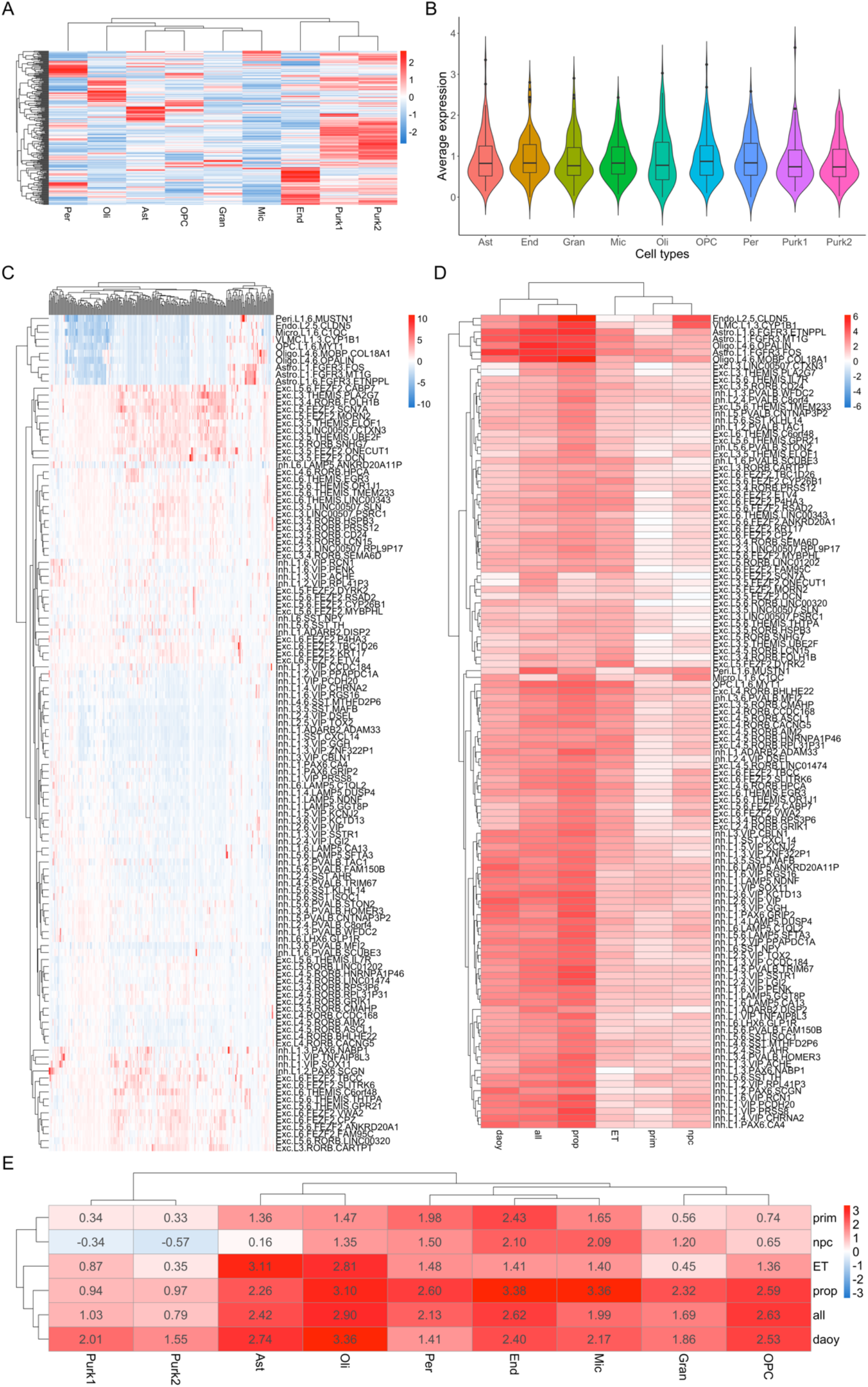
Single-cell enrichment of drug DEGs in cerebellar and cortical tissues. A. Single-cell enrichment Z-score heatmap of convergent propranolol/primidone DEGs in adult cerebellar tissue. Rows represent DEGs; columns indicate cell types; legend color scheme is based on enrichment z-score direction. B. Violin plot of average expression per cerebellar cell type of convergent propranolol/primidone DEGs. C. Single-cell enrichment Z-score heatmap of convergent propranolol/primidone DEGs in adult cortical tissue. Rows represent DEGs; columns indicate cell types; legend color scheme is based on enrichment Z-score direction. D. Enrichment Z-score heatmap of DEGs gene-sets from different conditions (see below for abbreviations) in single-cell data from adult cortex. E. Enrichment Z-score heatmap of DEGs gene-sets from different conditions in single-nucleus sequencing data from adult cerebellar tissue Rows indicate condition gene-sets; columns indicate cerebellar cell-types. Abbreviations: ET, ET related-genes; prop, convergent propranolol DEGs in both cell types; prim, convergent primidone DEGs in both cell types; DAOY, convergent propranolol and primidone DEGs in DAOY cells only; NPC, convergent propranolol and primidone DEGs in NPCs only; all, convergent propranolol and primidone DEGs in both cell types.

## DISCUSSION

Understanding the cellular and molecular mechanisms behind drug treatments can inform on disease pathophysiology. In this study, we sought to investigate the transcriptomic effects of first line treatments for ET in cerebellar DAOY cells as well as NPCs, to gain insight on potential disease related genes. We found that propranolol and primidone affected expression of multiple genes related to movement disorders and ET. Notably, *TRAPPC11*, whose expression was previously shown to be altered in ET cerebellar cortex and is also involved in protein trafficking^26^. Other genes related to endosomal trafficking were found to be differentially expressed after propranolol treatment, such as *MYO1E* and *SYNJ1*. Convergent DEGs also displayed an enrichment of genes related to the ESCRT complex, known to be a pillar of endosomal trafficking in neurons. These findings potentially increase the likelihood of endosomal trafficking being altered in ET and possibly partly restored through transcriptomic effects of propranolol.

Axon guidance was previously associated with ET in several studies^2,11,26,34,35^. Bulk-RNA sequencing of cerebellar cortex and dentate nucleus of ET patients showed a significant enrichment of axon guidance genes^26^. Hallmark axon guidance genes such as *ROBO1* (z-score = 5.87, q-val = 1.88E-06) and *NEO1* (z-score = 4.01, q-val = 5.04E-03) were both found to have increased expression following drug treatment. NEO1 (and its paralog DCC), which binds netrin-1, is implicated in cell-cell adhesions, mostly between axons and oligodendrocytes, as well as cell-extracellular matrix adhesions. Netrin-1 also acts on dendrite arborisation, increasing connections in excitatory synapses^36^. Interestingly, NEO1 protein remains expressed in Purkinje cells of the adult cerebellum (GTEx V8). Thus, the post-developmental role of axon guidance signalling pathways is to maintain adhesions and important synaptic connections between cells. This might be an important process by which ET tremorolytic drugs diminish tremor. These findings on axon guidance are concordant with other Reactome/GO-terms found to be enriched amongst DEGs, most notably semaphorin interactions, cadherin binding, and actin cytoskeleton reorganization. Purkinje cell axons in ET patients have shown accumulations of disordered neurofilaments (‘axonal torpedoes’) leading to abnormal axonal morphologies^35^. This process is thought to either be part of a neurodegenerative cascade or a response to neurodegeneration. Moreover, decreased neuronal density was observed in multiple brain regions of ET patients, most notably the inferior cerebellar peduncles through which afferent axons from the brainstem nuclei pass in order to reach the cerebellar cortex^37^. Our findings therefore provide additional support for the involvement of axon guidance molecules in ET pathophysiology.

We also identified the CaMKK2 signalling pathway as significantly enriched in propranolol DEGs in DAOYs and NPCs. CaMKK2 exacerbates Aβ42 synaptotoxicity in Alzheimer’s disease through Tau protein phosphorylation by AMPK^33^. This pathway is sensitive to cellular calcium intake, which was shown to be affected at the transcriptome level by both propranolol and primidone. Both Tau protein and amyloid-beta abnormalities have been observed in ET cerebellar tissues, with multiple findings pointing towards protein aggregation being a hallmark of the disease^38,39^. Propranolol affecting transcription of genes implicated in both CAMKK2 and Ca^2+^ signalling pathways might imply that ET drugs could reduce aggregate-induced neurotoxicity.

Convergent drug DEGs did not correlate with transcriptomic effects of common ET variants (TWAS DEGs). Moreover, propranolol and primidone DEGs displayed weak non-significant correlations with gene expression in the cerebellum of ET patients, the principal brain region affected in this disorder^1^. There are several possible explanations for these results. The relatively underpowered state (for a common disease) of the current ET GWAS might not capture the effects of common variation on transcription, in part explaining the absence of correlation with drug DEGs. Moreover, the lack of good cell models for cerebellar neurons as well as the neurodevelopmental state of NPCs also impair adequate comparisons between TWAS statistics and drug DEGs presented in this study.

Convergent drug DEGs are significantly more likely to be genes predicted to be intolerant to LoF variants. Mutationally constrained genes are more likely to be essential for cell homeostasis and survival and thus more likely to be implicated in disease when affected by LoF mutations^24^. Given that both ET drugs converged on these genes in multiple cell types increases the likelihood that these genes harbour rare variants associated with ET. Upregulated DEGs were found to be significantly less tolerant than all protein coding genes while downregulated DEGs were as tolerant as all protein coding genes. These genes could be good candidates for future targeted sequencing, especially within propranolol and primidone responsive cohorts.

Identifying cell types affected in ET remains difficult. Several conflicting studies have tried to identify specific pathological morphologies in post-mortem cerebellum of ET patients, most notably in Purkinje cells, yet no defining histopathological markers have been found^35^. Here we sought to identify the relevant ET cell types by assessing the enrichment of variant-harbouring ET genes within single cells in cerebellar and cortical tissues. Expression of ET genes were mostly enriched within L3-L5 excitatory neurons in the cerebral cortex, more specifically *FEZF2* L5 glutamatergic pyramidal neurons^40^. These neurons originate in the primary motor cortex (M1) and form the corticospinal tract that projects to lower motor neurons, which controls conscious movements. These neurons are influenced by multiple cortico-cortical pathways but also input from the cerebellothalamic tract, crucial for movement coordination. The primary motor cortex has previously been shown to be important for tremor generation in ET as subdural stimulation of M1 can reduce tremor intensity in patients^41^. Moreover, propranolol-targeted genes were mostly enriched in VIP-expressing inhibitory neurons of L1-L3. These neurons are known to inhibit motor neurons through different cortical pathways^42^. The enrichment of ET genes within M1 pyramidal neurons coupled with the enrichment of ET drug genes in motor neuron-inhibiting cells does suggest new potential cellular mechanisms through which tremor generation (and/or reduction) occurs in ET.

In the cerebellum, both ET genes and convergent drug DEGs were significantly enriched within astrocytes in the cerebellum. This somewhat contradicts previous histopathological findings postulating that Purkinje cells were the defining cell type in ET pathophysiology. Not much is known about the role of astrocytes in ET but based on other neurodegenerative diseases, it could be argued that they may play an important role in the onset or development of the disease^35^. Oligodendrocytes, whose dysfunction contributes to numerous other neurological diseases, also showed an enrichment of propranolol and primidone-targeted genes. Both astrocytes and oligodendrocytes might be targeted by ET drugs to reduce tremor since non-neuronal cell types are known to be involved in neurodegeneration in numerous diseases^43^. The lack of single-cell data on ET tissues is a limitation in the study of this disease but our results highlight a possible role for non-neuronal cells in the cerebellum in ET.

This study has a number of limitations. Propranolol and primidone are known to act on cell excitability and this effect was postulated as being important for tremor reduction in ET. Given that DAOYs and NPCs are non-excitable, it is very hard to assess the electrophysiological effects of these drugs in these cells. Moreover, the electrophysiological effects of drugs on cells are known to influence transcription^44^. This might explain why primidone had such a mild effect on transcription in both DAOYs and NPCs. Cells used in this study do not represent the complete range of of cell types in the cortex and cerebellum. NPCs do not completely replicate neuronal expression and do have a more neurodevelopmental transcriptomic state. DAOYs, on the other hand, are derived from cancerous cells and do have dysregulated expression of genes related to cell division and cell growth. Nevertheless, this study only serves as an ET drug effect screen and remains a steppingstone for more in-depth studies.

Our study identifies multiple cellular and molecular pathways implicated in ET pathophysiology and tremor reduction by both propranolol and primidone. Our findings also suggest a role for genes harbouring potentially rare, deleterious variants associated with ET. Targeted sequencing of these convergent drug genes in case-control cohorts could help to confirm or infirm this hypothesis. These genes could also be used as biomarkers for propranolol treatment in responsive ET patients. Our results also identify several cell types involved in ET in both cerebellar and cortical tissues. We also identify cell types potentially affected by propranolol and primidone through which tremor might be reduced in ET. Future studies will be needed to further identify the transcriptomic and electrophysiological effects of both drugs, possibly using more representative neuronal models such as iPSC-derived Purkinje cells, non-neuronal cell types as well as motor neurons. Moreover, single-cell experiments studying the transcriptomic effects of ET drugs on patient-derived tissues will be required to understand the complex nature of this disease.

---

**Supplementary Figure 1.**
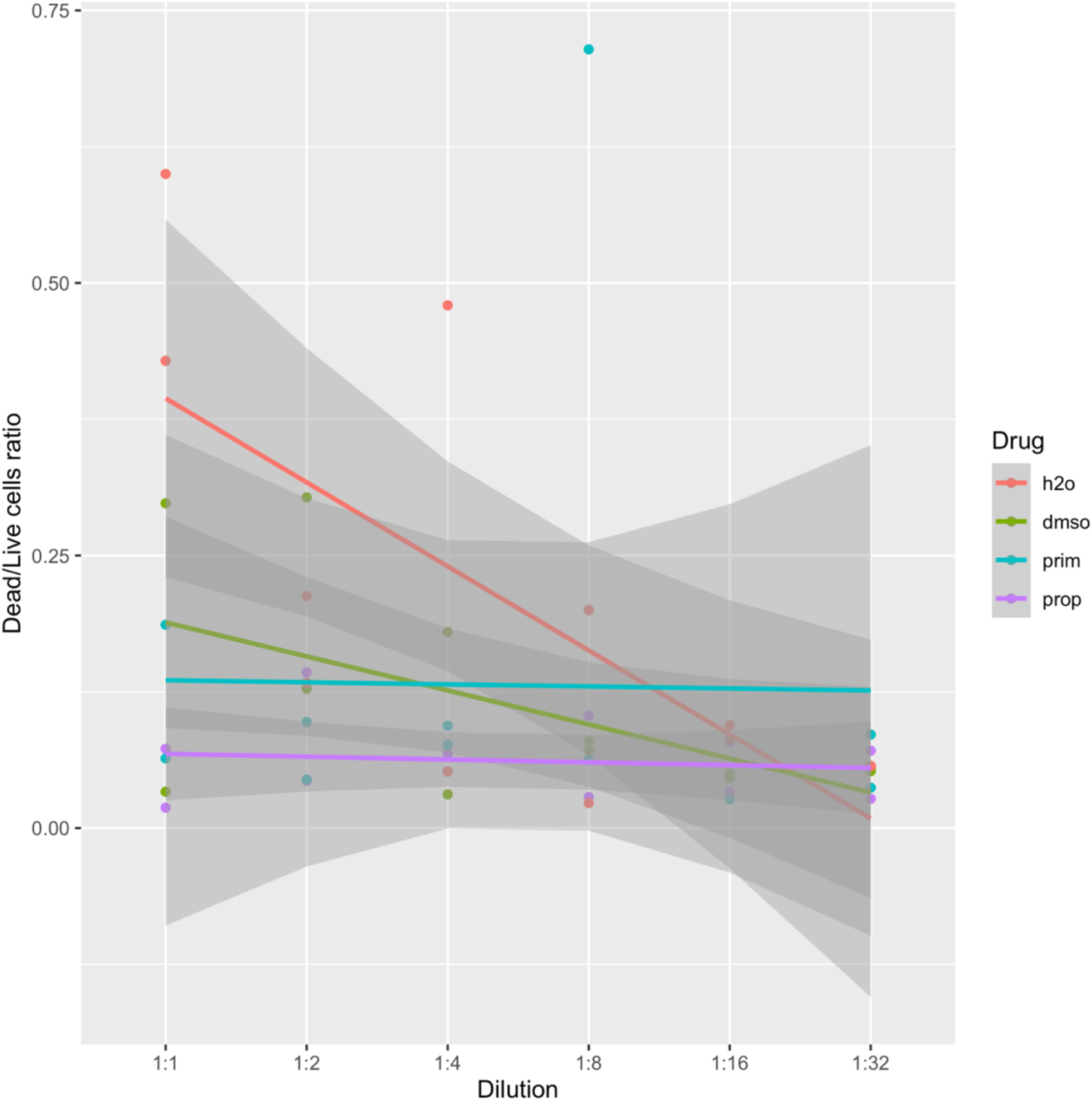
DAOY kill curve. Dead over live cell ratios were calculated based on NucGreen and NucBlue (DAPI) staining after 5 days of treatment. Dilutions are calculated from initial concentrations of drugs or DMSO (%; corresponding to the percentage of DMSO that primidone was diluted in). 1:1 dilutions; Propranolol = 0.0156 μg/mL, Primidone = 25 μg/mL; DMSO = 0.235%.

**Supplementary Figure 2.**
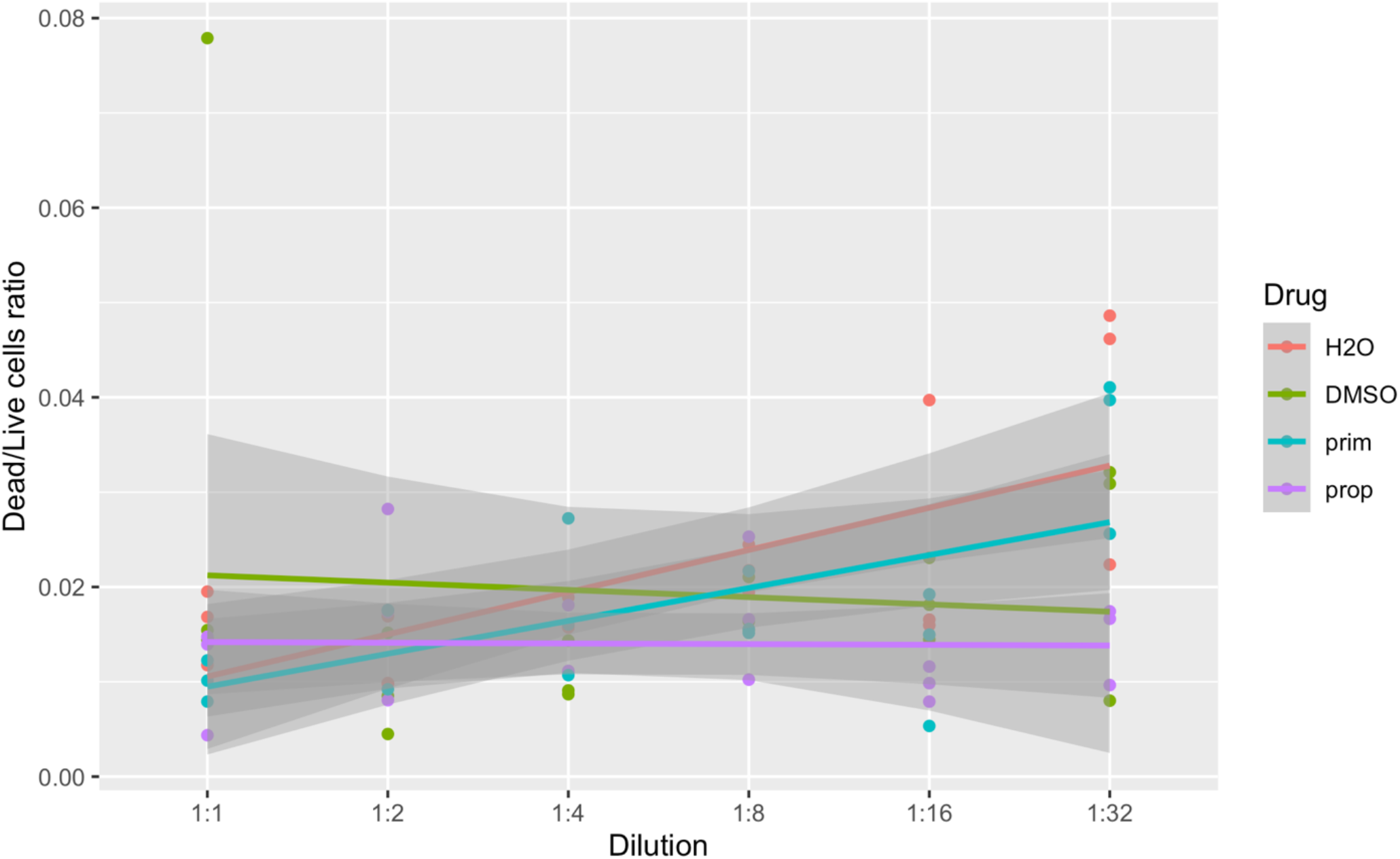
NPC kill curve. Dead over live cell ratios were calculated based on NucGreen and NucBlue (DAPI) staining after 5 days of treatment. Dilutions are calculated from initial concentrations of drugs or DMSO (%; corresponding to the percentage of DMSO that primidone was diluted in). 1:1 dilutions; Propranolol = 0.0156 μg/mL, Primidone = 25 μg/mL; DMSO = 0.235%.

**Supplementary Figure 3.**
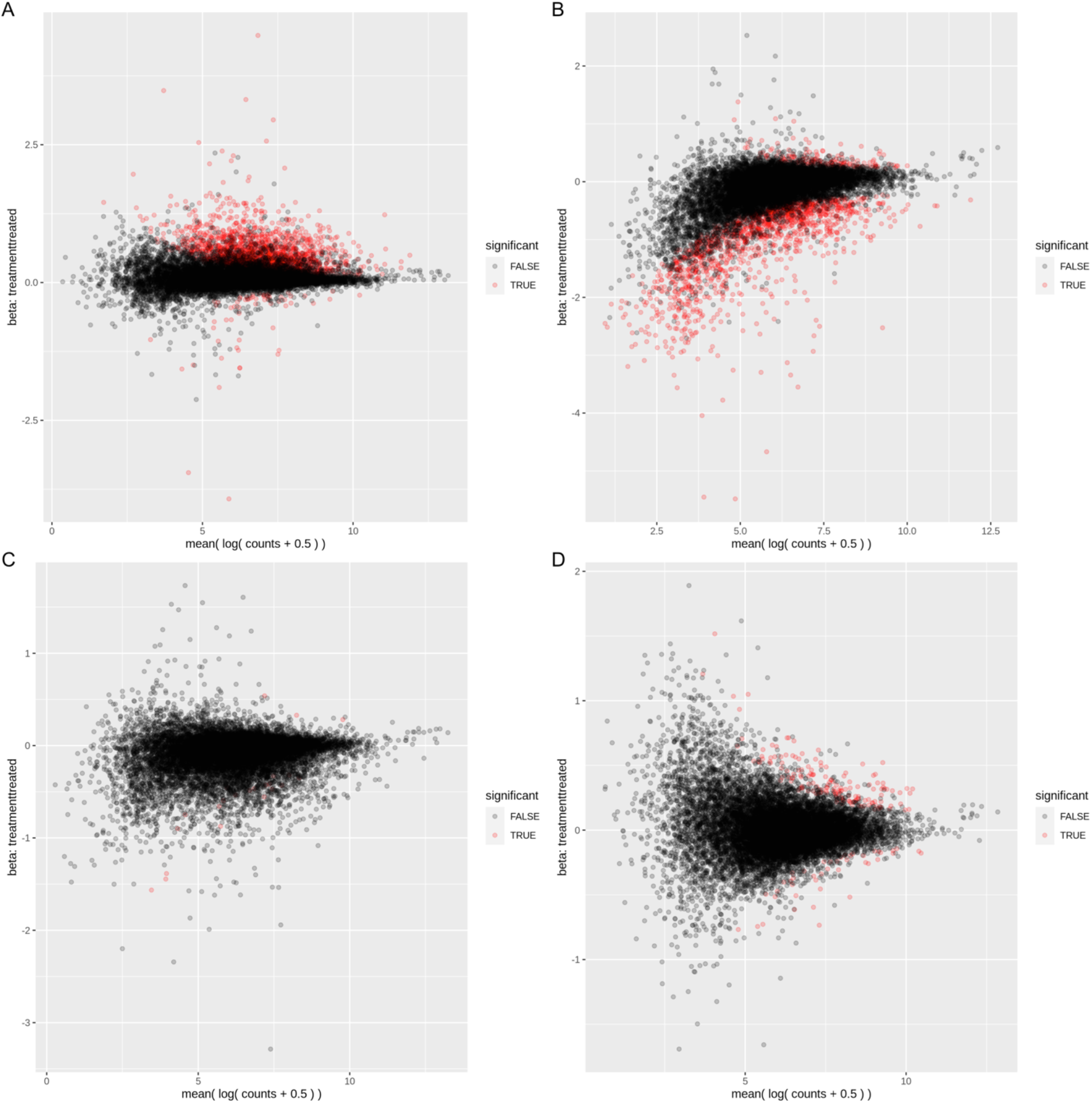
Mean A plots. A. DAOYs treated with propranolol. B. DAOYs treated with primidone. C. NPCs treated with propranolol. D. NPCs treated with primidone

**Supplementary Figure 4.**
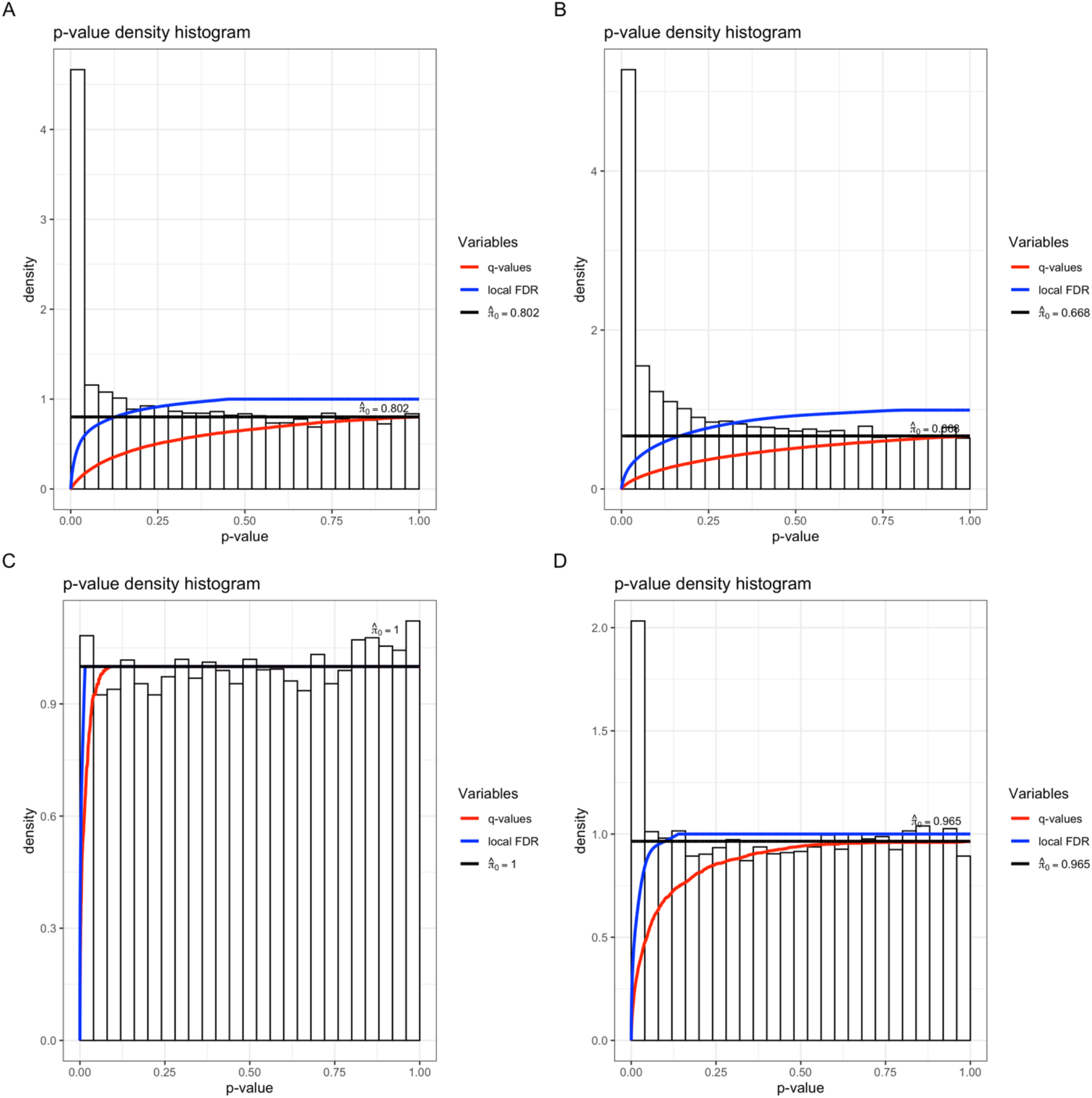
P-value histograms. A. DAOYs treated with propranolol. B. DAOYs treated with primidone. C. NPCs treated with propranolol. D. NPCs treated with primidone.

